# Disrupted chromatin architecture in olfactory sensory neurons: A missing link from COVID-19 infection to anosmia

**DOI:** 10.1101/2022.08.19.504545

**Authors:** Zhen Wah Tan, Ping Jing Toong, Enrico Guarnera, Igor N. Berezovsky

## Abstract

We tackle here genomic mechanisms of a rapid onset and recovery from anosmia - a useful diagnostic indicator for early-stage COVID-19 infection. On the basis of earlier observed specifics of olfactory receptors (ORs) regulation in the mice chromatin structures, we hypothesized that the disruption of OR function can be caused by chromatin reorganization taking place upon SARS-CoV-2 infection. We reconstructed the chromatin ensembles of ORs obtained from COVID-19 patients and control samples using our original computational framework for the whole-genome chromatin ensemble 3D reconstruction. We have also developed here a new procedure for the analysis of fine structural hierarchy in local, megabase scale, parts of chromosomes containing the OR genes and corresponding epigenetic factors. We observed structural modifications in COVID-19 patients on different levels of chromatin organization, from alteration of the whole genome structure and chromosomal intermingling to reorganization of contacts between the chromatin loops at the level of topologically associating domains. While complementary data on known regulatory elements point to pathology-associated changes within the overall picture of chromatin alterations, further investigation using additional epigenetic factors mapped on 3D reconstructions with improved resolution will be required for better understanding of anosmia caused by SARS-CoV-2 infection.

## Introduction

The COVID-19 pandemic, caused by the SARS-CoV-2 coronavirus, has placed significant strain on medical resources worldwide, and early detection and isolation has been the primary strategy by which governments worked to contain the disease spread. To enable early detection, surveys on early symptoms of SARS-CoV-2 infection uncovered anosmia as a useful predictor of COVID-19 infection (1,2), showing higher positive and negative predictive values than other flu-like symptoms (3,4). However, the mechanisms behind the rapid onset of anosmia has remained controversial, as a large proportion of COVID-19 patients with anosmia did not experience nasal congestion (5,6), the causes of anosmia, reasons behind impaired olfaction extend beyond physical obstruction of odorants in the olfactory epithelium (OE). The SARS-CoV-2 virus enters human host cells via a mechanism similar to the SARS-CoV virus: the proteolytic cleavage of the viral spike protein by TMPRSS2 and entry initiation by binding to the host ACE2 receptor (7). Since olfactory sensory neurons (OSNs) do not express ACE2 and TMPRSS2 at significant levels, it has been proposed that viral infection of non-neuronal cell types in the olfactory epithelium (OE) are the main factors behind impaired olfaction (8). Among these, sustentacular (SUS) cells — which provide structural support in the OE — express high levels of both viral receptors (8,9), and infection of SUS cells may impair metabolic and structural functions that are critical for proper OSN functioning (10,11). However, these changes are not likely so drastic as to lead to OSN cell death, as most anosmic patients recover their sense of smell within 1-2 weeks (12–15), shorter than the time required for OSN replacement and maturation, and axon/cilia growth (16–18). On the other hand, inflammatory response of the innate immune system in the OE may also contribute to impaired olfaction, as several studies detected high levels of the pro-inflammatory cytokines TNF-α and IL-6 in the OE of SARS-CoV-2 infected patients (19,20).

While many studies linked released cytokines to changes in the cellular microenvironment that can interfere with OSN function or induce premature apoptosis (19,21), a few have focused on how inflammatory response in the OE may affect olfactory function at the level of gene expression. For example, in a mouse study, sterile induction of innate immune signaling in the OE was associated with diminished odor discrimination (22). The OSNs showed significantly reduced expression of olfactory receptors genes (ORs) — these encode G protein-coupled receptors that bind to odorants and trigger olfactory signal transduction — suggesting that olfactory dysfunction in response to inflammation may be traced to altered OR regulation in OSNs.

The human OR repertoire consists of 376 functional genes distributed across 18 chromosomes, and can be categorized into distinct phylogenetic groups, Class I and Class II, by sequence homology. Most ORs are located in clusters up to ~1Mbp in size, and it has been widely accepted that in the maturation process of OSNs, the neurons begin to exhibit specialized OR expression, and each mature OSN (mOSNs) expresses only one allele of one randomly selected OR gene (23,24). With gene translocation ruled out as a possible mechanism for OR selection (25), current models based on mouse studies propose that the control of mOSN OR expression is achieved via multiple levels of chromatin organization. At the whole-genome level, immuno-fluorescence studies have found that olfactory sensory neurons exhibit an inverted nuclear architecture: While typical cells are organized with constitutive heterochromatin (cHC) localized the nuclear lamina, OSNs show an aggregation of a large cHC block away from the lamina, surrounded by facultative heterochromatin (fHC) (26). The observed tendency for ORs to colocalize in the cHC or fHC regions may facilitate the selection of a single gene allele for expression (27,28). This inverted architecture appears to also be mediated by the downregulation of lamin-B receptors (LBR) in OSNs, as restoring LBR expression led to a decondensation of the cHC block, and a concomitant disruption of OR aggregation and expression (27). At a finer structural scale, epigenomics and reporter assays have also identified a set of enhancers located in the vicinity of mouse ORs (termed the Greek Islands) (29): these regulatory elements were observed to physically interact with activated ORs and with one another, even across different chromosomes (30,31). In view of these two levels of chromatin organization, a plausible picture of OR regulation in OSNs may consist of the aggregation of OR genes in the HC block, enabling the selection of one OR gene allele to be decompacted and activated through a looping interaction with an interchromosomal enhancer hub (31).

To understand how SARS-CoV-2 infection may lead to anosmia through OR dysregulation and chromatin structure disruption, we began by using publicly available Hi-C data of OSNs of SARS-CoV-2-infected and control human subjects (20), obtaining the whole-genome reconstructions of the chromatin ensemble in each subject (32). Focusing on the detailed organization of OR clusters, we then analyzed Hi-C data to identify fine structural units of chromatin and potential functional links between them. We adopted a two-pronged approach in characterizing chromatin structural differences in SARS-CoV-2-infected patients and controls, on one hand studying large-scale structural shifts by spatial reconstructions at the genome level, and on the other hand zooming in to fine-scale organization of key gene clusters. Firstly, we use a Markov state model to identify coarse Mb-scale structural units (33), then apply a stochastic embedding procedure (32) to obtain genome-level reconstructions of the chromatin ensemble. Secondly, to understand chromatin organization and packing at the local level we devised a novel method for identifying localized structural units at the ~30 to 200-500 kbp scale, mapping out how strong links between these units shape physical packing of the chromatin fiber from chromatin loops (34–36) to (sub)TADs (37). Olfactory regulation is recognized as an involving inter-chromosomal processes in mice (31), with olfactory receptors organized in multiple gene clusters spread across multiple chromosomes in both mice and humans. This work provides a first look at how normal human OSNs exhibit similar features of chromatin organization to that observed in studies of normal mouse OSNs, highlighting how the disruption of OSN chromatin structure may be an important link in the development of anosmia in SARS-CoV-2 patients.

## Materials and Methods

### Reconstruction of 3D chromatin ensemble structure

We investigate here large-scale changes in chromatin structure captured in Hi-C chromatin interaction data for OSNs from 4 SARS-CoV-2-infected patients and 2 control subjects, by reconstructing of the chromatin ensemble structure (Figure 1 and Supplementary File, Figure S1). Our approach consists of two steps, by first (i) identifying megabase-level structural partitions using a Markov state model (MSM) of Hi-C interactions (33), and second (ii) obtaining ensemble reconstructions at the level of these partitions using a stochastic embedding procedure (SEP) (32), as implemented in software pipelines published previously.

**Figure 1.**
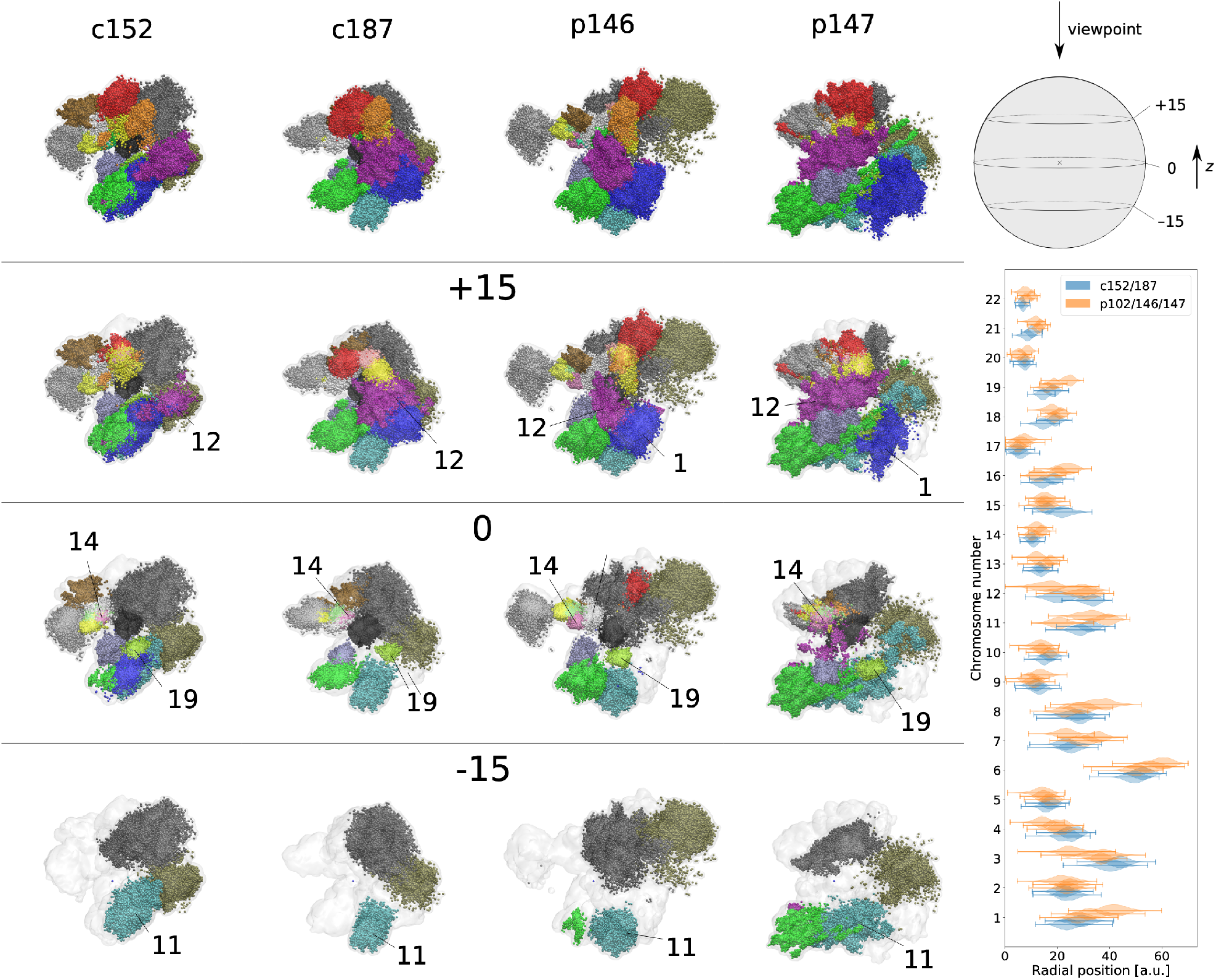
Reconstructions of whole-genome chromatin ensemble for olfactory sensory neurons in SARS-CoV-2-infected patients and controls. Left, top row: Full reconstructions colored by chromosomes are shown for controls (c152, c187) and patients (p102, p146, p147). (bottom left) Subsequent rows below are cross-sectional views of the respective reconstructions, at the positions indicated by the schematic (top right), where *z* = 0 intersects the centroid. The approximate centers of chromosomal territories 1, 9, 11, 12, 14, and 19 are indicated on the cross sections. Radial distributions of chromosomal territories, comparing the same chromosomes on control (blue) and patient (orange) samples.

In the first step, we partition the chromatin interaction network using a MSM approach. By modelling the diffusion of small molecules in densely packed chromatin, the MSM enables one to partition the genome into structural units where epigenetic factors tend to be localized, which provides a suitable level of coarse-graining of genome structure as captured by Hi-C. For each sample, we identified more than 1500 structural units with average size 2Mbp across the genome. To then obtain spatial embeddings of chromatin ensemble structure, we define a matrix of effective interactions between the structural partitions by coarse-graining the Hi-C matrix. These effective interactions are subject to stochastic fluctuations in the cell population, and we applied the SEP method to reconstruct the chromatin ensemble through sampling of structural variations between and within individual structural units.

We identified structural units in the genome by applying a MSM partitioning procedure (33) to Hi-C data using a metastability criterion of *ρ* < 0.9, yielding 950, 901, 850, 952, 800, and 903 structural units for samples c152, c187, p102, p116, p146, and p147, respectively. Effective interactions between the structural units are defined by coarse-graining the Hi-C matrices. To map effective interaction matrices to ensemble structures, we used the Stochastic Embedding Procedure we developed previously (32), via a distance geometry approach for spatial embedding of effective interaction data subject to stochastic fluctuations. We obtained *N* = 100 samples of fluctuations in effective interactions using a gaussian model with strength factor *α* = 0.2. The sampled interaction matrices were normalized using the SCPNs algorithm described previously (32), before using spatial embedding to obtain positions of individual structural units. We then modelled each structural partition as a spherical crumpled globule with radius *r* scaling with partition size *s*, *r* ~ *s*^0.6^, representing intra-partition structural variability by random sampling of points within the partitions at 1Mbp intervals. The merged samples represent regions of space accessible to individual partitions in the ensemble of chromatin structural states. Without information on the gender of subjects, we have focused structural analysis only on somatic chromosomes 1-22 in this work.

### Robustness of chromatin structural variation between samples

To evaluate the robustness of ensemble reconstructions and the significance of changes in chromatin structure, we repeated the 2-step reconstruction procedure for each Hi-C sample by performing 3 independent MSM partitioning runs (MSM replicates), followed by 2 independent SEP ensemble samplings for each obtained partitioning (SEP replicates). The 6 replicate reconstructions per Hi-C sample are shown in Supplementary File, Supplementary File, Figure S2 along with the network of chromosomal intermingling, and we show a quantitative comparison of these ensemble reconstructions in Supplementary File, Supplementary File, Figure S3. The bottom-left of the latter shows nuanced differences between reconstructions of the same Hi-C sample reflected in shifts in the distance correlation functions g(r), while the top-right shows the large-scale changes in chromosomal mixing fractions between all reconstructions.

Firstly, as distance correlations *g*(*r*) capture more detailed information on chromosome morphology, we used the histogram difference Δ*g*(*r*) (see Materials and Methods) to quantify more nuanced structural shifts between reconstructions of the same Hi-C sample (see bottom left of Supplementary File, Figure S3). We found essentially no structural difference between SEP replicates, indicating that the sampling of interaction fluctuations is sufficient to yield robust ensemble representations. Between MSM replicates of the same Hi-C datasets (except for p146), we observed only weak differences in *g*(*r*) owing to minor shifts in chromosomal positioning. For p146, visual inspection of the structures and g(r) correlation functions of MSM replicates showed that the reconstructions maintained consistent relative positioning of chromosomal territories, despite slightly larger variation in bulk distances between non-intermingling chromosomes that led to the larger histogram differences Δ*g*(*r*) seen in Supplementary File, Supplementary File, Figure S3. Secondly, to study large-scale structural differences between samples, we focused on changes in chromosomal intermingling and quantified the mixing fraction *m_ij_* of chromosome *i* in chromosome *j* as the fraction of chromosome *i* located within a cutoff distance 1 a.u. from chromosome *j* (see Materials and Methods). Comparing all 231 chromosome pairs across all 36 ensemble reconstructions, 69 chromosomal pairs consistently showed no observable intermingling. Among the remaining 162 chromosomal pairs, Supplementary File, Figure S3 shows that mixing fractions were largely consistent between MSM and SEP replicates of each Hi-C sample, compared to the differences between Hi-C samples. While some differences exist between the 2 control samples, we observed a higher degree of heterogeneity between the 4 patient samples (Kruskal-Wallis H-test on the first principal components of control and patient samples, yielding p-values 4 × 10^-3^ and 2 × 10^-4^), with p147 and p116 showing greater and less chromosomal intermingling respectively. Importantly, Supplementary File, Figure S3 shows significant differences between controls and patients in chromosomal intermingling, with several chromosome pairs showing strong shifts between control and patient chromatin states (38 chromosomal pairs with Mann-Whitney U-test p-value < 0.1%). The ensemble reconstruction procedure implemented was thus able to capture structural variation within the control and patient sample groups, and identify consistent structural changes in OSNs for SARS-CoV-2-infected patients.

### Quantifying intermingling between chromosomal territories

We quantified the degree to which chromosomal territories intermingle using ensemble reconstructions obtained from the stochastic embedding procedure. To that end, we defined the mixing fraction *m_ij_* from chromosome *i* to chromosome *j* as the fraction of sampled points in *i* within a cutoff distance of 1.0 a.u. (corresponding to a physical distance of approximately 350nm) from sampled points in *j.* In this work, each chromosome is represented by 5000 to 25000 points (proportional to chromosome size), and a mixing fraction of 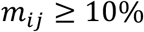 constitutes significant intermingling. For two ensemble reconstructions *r*_1_ and *r*_2_, we quantified the difference between mixing fractions by the simple difference 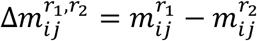.

### Distance correlation functions g(r) of single chromosomes or chromosome pairs

To capture more detailed information on chromosomal morphology, we determined the pairwise distance correlation functions *g*(*r*) for chromosome pairs, as well as for individual chromosomes (Figure 2 and Supplementary File, Figure S4). The distance correlation *g_ij_*(*r*) for chromosomes *i* and *j* is defined as the probability density that two randomly sampled points in the chromosome pair is separated by a distance *r*: from ensemble reconstructions, we computed the *g*(*r*) functions as a normalized histogram of sampled pairwise distances. The single-chromosome correlation functions *g_i_*(*r*) are defined and computed similarly, considering only sampled points in one chromosome. The difference between distance correlations in two ensemble reconstructions *r*_1_ and *r*_2_ is quantified by the histogram distance:

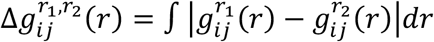

**Figure 2.**
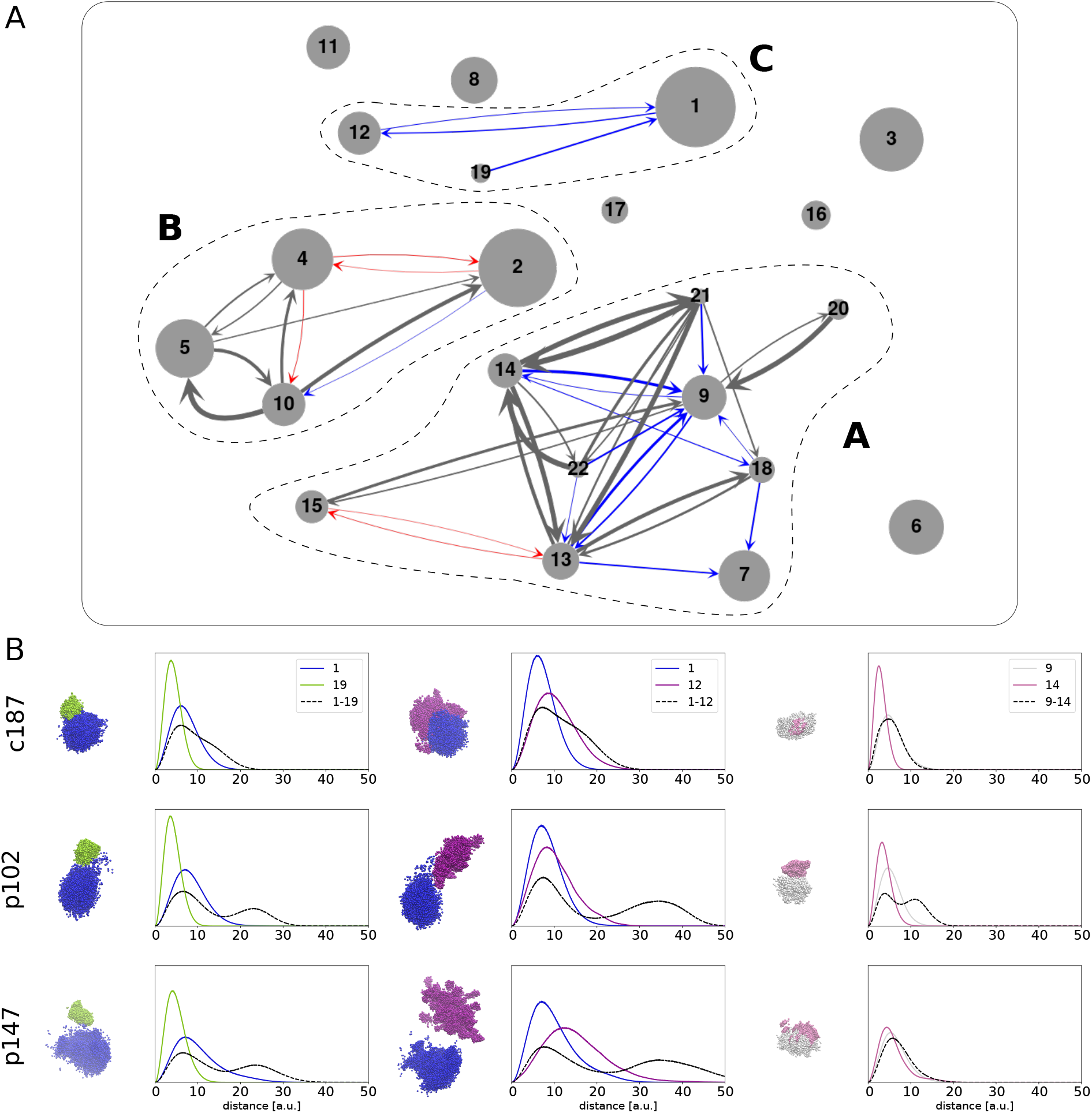
Differences in chromosomal intermingling between control and patient samples. **A.** Chromosomal intermingling network in ensemble reconstructions. Chromosomes are represented as nodes with radius scaling with chromosome size. Edges connect chromosomal pairs that intermingle strongly in all controls only (blue), all patients only (red), or in all controls and patients (grey); edge width represents the average mixing fraction observed in corresponding samples. Three chromosomal groups can be identified in the network, as marked by dashed lines. **B.** Typical chromosomal morphology and distance correlation functions *g*(*r*) of chromosome pairs 1-19, 1-12, and 9-14 in controls and patients. Chromosomes adopt the same color convention as in Figure 1: 1 – blue, 9 – white, 12 – purple, 14 – pink, 19 – lime green.

### Human olfactory receptors, OSN markers, and putative enhancers mapped from mouse data

We retrieved a list of identified human olfactory receptors (ORs) from the HORDE database (38), which contained a total of 376 functioning genes, after removing 439 pseudogenes. Genomic coordinates for each gene were then identified on the hg38 human reference genome. The OR class is identified by the gene family ID: ID numbers 51 to 56 are categorized as Class I, while all others belong to Class II. A list of 15 marker genes for mature human OSNs was retrieved from (24). A list of 35 mouse olfactory enhancers, termed Greek Islands, was identified by the Lomvardas Lab (29). We mapped the genomic coordinates of these enhancers to the human reference genome by successive application of the UCSC LiftOver tool (https://genome.ucsc.edu/cgi-bin/hgLiftOver), from mm9 to mm10, then from mm10 to hg38. Among the 25 successfully mapped regions, we further required that putative enhancers (PEs) be located at most 1Mbp from the nearest human OR, yielding a final list of 15 PEs. The full list of mouse enhancers’ mapped genomic coordinates, and proximity to ORs are provided in Supplementary File, Table S1.

### Local structural unit analysis of fine chromatin structure

To understand how chromatin is structured and packaged at a local level, we developed a bottom-up approach to identify localized structural units beginning with intra-chromosomal Hi-C data at 5kbp resolution. We quantify the degree of association between two structural units *i* and *j* by the ratio *r_ij_* of the interaction energy (given by total Hi-C interaction counts) between units *i* and *j*, denoted *F_ij_*, to the geometric average of interaction energy within the individual units (ignoring self-interactions at the 5kb level), denoted *F_i_*, *F_j_* respectively: 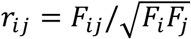. The procedure begins by considering each 5kb bin as a separate structural unit, and iteratively merges pairs of structural units with the highest value of *r_ij_*. The global maximum value of *r_ij_*, denoted *r_max_*, decreases monotonically with each iteration, and at lowest *r_max_* = 1.0 the structural units of first level of hierarchy are obtained. Continuing with the iterative procedure reduces *r_max_*, yielding larger structural units that are more physically separated at lowest *r_max_* = [0.8, 1), [0.6, 0.8), and [0.5, 0.6), revealing the hierarchical organization of chromatin structure. The result of applying the procedure on a 1Mb region is represented schematically in Figure 3 (see also Supplementary File Figure S), where the top shows 1D representations of the structural units at different levels, and the bottom shows how individual 5kb bins (black nodes) are grouped into structural units at different levels of hierarchy (grey, blue, red, and green ellipses). Supplementary File, Table S2 lists the average unit size at lowest *r_max_* = 1.0,0.8,0.6,0.5 for chromosomes 1, 11, and 14 in each of the samples c152, c187, p102, and p146. For the sake of definition from now on, the *r_max_* = 1.0, 0.8, 0.6, 0.5 designate *r_max_* ≥ 1, [0.8, 1), [0.6, 0.8), and [0.5, 0.6), respectively.

**Figure 3.**
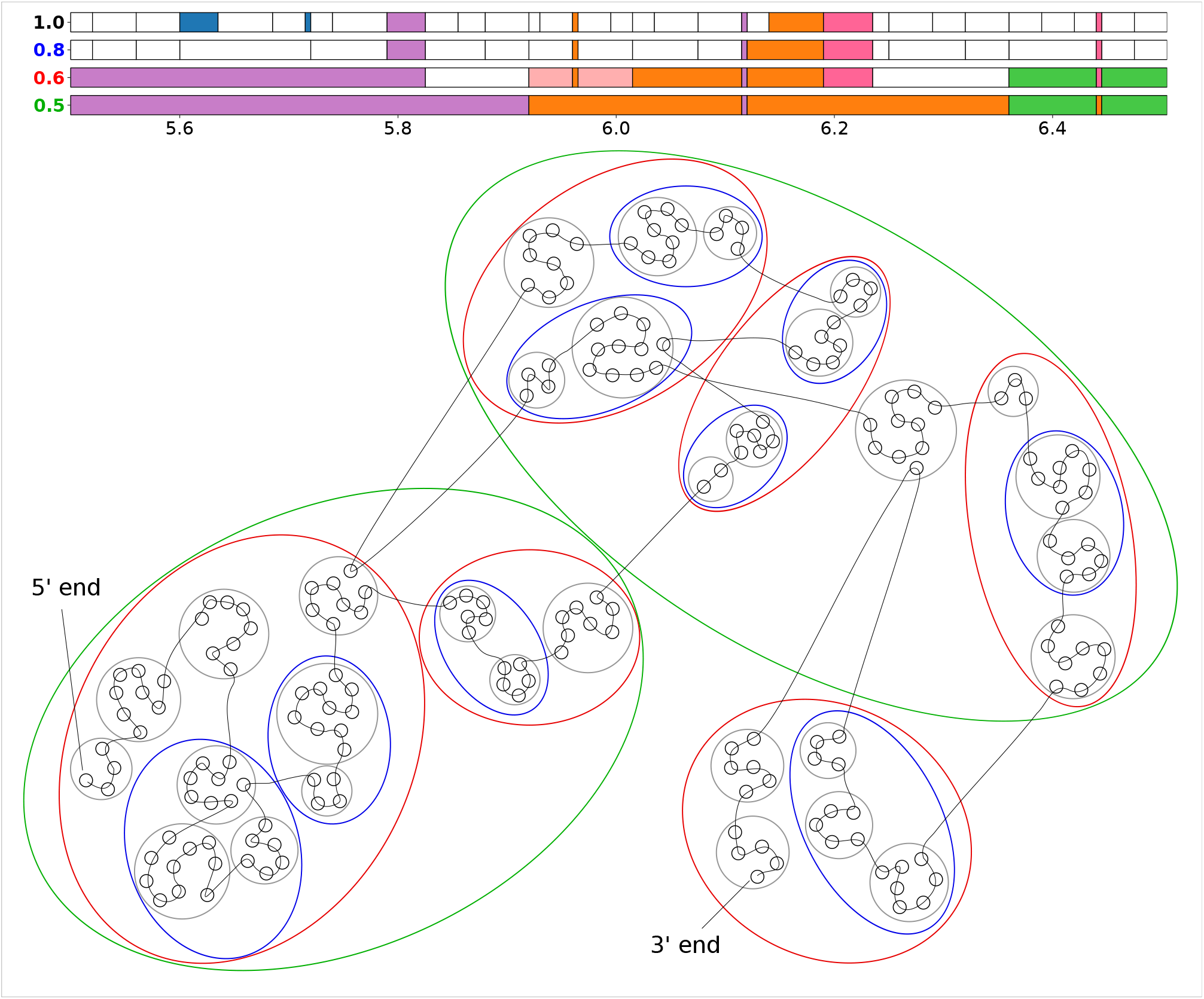
Iterative procedure for identifying fine structural hierarchy in chromosomes. This figure shows structural units identified at various levels of hierarchy in a 1Mb region (5.5-6.5Mb) of chromosome 11. In the network representation, individual 5kb bins/loci in this region are represented by black circular nodes, with a chromosomal trace overlaid to show the relative genomic locations of each locus. Beginning by considering each locus as an individual structural unit, we iteratively join pairs of structural units with the highest interacting energy ratios *r*, to a point where the largest energy ratio *r_max_* falls below 1.0, yielding structural units denoted by the grey circles. Continuing with this procedure, we obtain different levels of hierarchy when *r_max_* falls below 0.8, 0.6, and 0.5, which give structural units represented by the blue, red, and green ellipses, respectively. 1D representations of the structural units (in the top) obtained at each level of hierarchy are denoted by the value of *r_max_* on the left. Units composed of a single contiguous genomic segment are shown as white horizontal bars in the plot, while units composed of non-contiguous segments are color-coded accordingly: e.g., the two blue segments and the two orange segments at *r_max_* = 1.0 form two corresponding discontinuous structural units.

## Results

### Reorganization of chromosomal territories in OSNs of SARS-CoV-2-infected patients

We use here recently developed two-step procedure consisting of Markov state modelling for identifying structural units and stochastic embedding procedure for obtaining the 3D ensemble reconstruction of the whole-genome chromatin structure, elucidating morphology of chromosomal territories (CTs) and intermingling between them. Inspecting obtained ensemble reconstructions of OSN chromatin visually (top rows in Figure 1 and Supplementary File, Figure S1) we found that although the overall organization of chromosomal territories (CTs) shows some degree of similarity across all samples, patient samples yield noticeable shifts and diversity in CT positioning, morphology and intermingling. Mouse studies have indicated that olfactory regulation in OSNs depends upon inter-chromosomal interactions between key regions (30,31): given that the human homologs of these genomic regions are also distributed across different chromosomes, it is likely that similar inter-chromosomal interactions may also be critical for OSN regulation in humans. One major class of OSN-associated genes are the olfactory receptor genes (ORs), which bind odorants in the nasal passage and enable odor perception and discrimination. ORs are distributed across 17 somatic chromosomes in humans, of which the largest sets are located on chromosomes 11 (166 ORs), 1 (62), 9 (25), 14 (23), 19 (20), and 12 (17) — the centers of these chromosomes are labelled in crosssectional views below the full reconstructions in Figure 1 (a larger, equally-spaced series of crosssectional views is given in Supplementary File, Figure S1). We observed that chromosomes 9, 14, and 19 were consistently located near the center, chromosomes 12 and 11 - near the periphery in all 5 samples, while chromosome 1 shifted towards the nuclear periphery in infected patients. Comparing the radial distribution of CTs (right panel of Figure 1) confirms that, indeed, while the radial positioning of most chromosomes remained unchanged between controls and patients, chromosome 1 was noticeably displaced outwards in OSNs of infected patients.

To systematically quantify and compare CT organization, we identified chromosome pairs that showed consistent and strong intermingling in either controls or patients. In this work, chromosome i is defined as strongly intermingling with chromosome *j* if the mixing fraction *m_ij_* exceeds a threshold value of 10% (see Materials and Methods). Our results are summarized by the network in Figure 2A, where grey arrows indicate strong intermingling present in all 5 samples, while blue/red arrows indicate strong intermingling in all control/patient samples, respectively (see Supplementary File, Figure S4 for separate intermingling networks for each sample). Two groups of intermingling CTs can be identified in all samples, demarcated by dashed outlines in Figure 2. Firstly, chromosomal group A comprises mainly of smaller chromosomes, with four of the acrocentric chromosomes (13, 14, 21, 22) showing exceptionally strong intermingling in both controls and patients. The data also showed chromosome 9 to be a central component of the structure of group A in controls, forming strong intermingling with almost all chromosomes in the group. However, many of these structural links were lost in the patient sample p102, resulting in a sparser network of CT intermingling in group A for patients. Secondly, chromosomal group B consists of larger chromosomes forming highly consistent and strong intermingling in all samples, with slightly stronger intermingling overall in patients. Control samples also formed a smaller, third chromosomal group C, where chromosomes 12 and 19 intermingled strongly with chromosome 1. The decoupling of chromosome 1 is especially evident in patient samples p147 and p102, due to the displacement of chromosome 1 towards the periphery as observed above.

Since many structural differences between controls and patients involved chromosomes with a large number of ORs, we sought to also understand detailed changes in CT morphology by studying how one-chromosome and two-chromosome distance correlation functions (see Figure 2B and Supplementary File, Figure S5) are different between controls and patients, focusing on c187, p102, and p147 as representative examples. The distance correlation function *g*(*r*) is a quantitative description of the morphology of non-rigid structures, and is defined as the probability distribution of distances between any two randomly selected points in the structure. Typically, CTs form an approximately globular structure that has the distance correlation *g*(*r*) forming a single peak at a position corresponding to the size of the globule: for example, in c187, chromosomes 1 and 19 have sizes of about 8 and 4 a.u., respectively (Figure 2B). Long tails in the distribution signal a deviation from sphericity, typically due to diffuse boundaries or elongated protrusions from the structure, such as the case of chromosome 1 in p102 and p147 (Figure 2B). Two-chromosome distance correlations can exhibit a range of functional forms depending on the degree of intermingling between the chromosomes. On one end of the spectrum, for chromosome pair 1-19 in p147 we observed a well-separated double peak structure in *g*(*r*), which indicates a clear separation of chromosomal territories — the first peak at 8 a.u. is the average size of individual CTs, and the second peak at 25 a.u. indicates the separation between the CT centers. Towards the other end of the spectrum, in c187 we observed that the *g*(*r*) curve for chromosome pair 1-19 contains only one dominant peak at 8 a.u., indicating strong intermingling between the two CTs. Comparing the structures and *g*(*r*) plots for chromosome pairs 1-12 and 1-19, it is evident how the consistent, strong intermingling between these chromosome pairs in control OSNs (exemplified by c187 in Figure 2B) was weakened or lost in infected patients (e.g., p102 and p147 in Figure 2B), potentially impacting processes that depend on inter-chromosomal interactions within chromosomal group C. Turning to chromosomal group A, where we observed the weakened intermingling of chromosome 9 with many other CTs in the group, we focused on the chromosome pair 9-14, both of which contain a large number of ORs. In c187, we observed that the two-chromosome *g*(*r*) is virtually indistinguishable from the one-chromosome *g*(*r*) for the larger chromosome 9, indicating exceptionally strong intermingling between these CTs. Reconstructions of patient OSNs, however, showed more diversity in CT morphology and intermingling. In p102, the distinct double peak structure in *g*(*r*) indicates a clear separation of CTs with minimal intermingling despite having a shared boundary. On the other hand, in p147, chromosome 14 formed a more dispersed CT that overlaps significantly with chromosome 9. The structure of chromosome 14 in p146 (see Supplementary File, Figure S3) represents an intermediate state between p102 and p147, where a poorly resolved secondary peak in pair-wise *g*(*r*) indicates a moderate degree of intermingling between the territories of chromosomes 9 and 14. Several cases of chromosomal intermingling consistently seen in OSNs of control subjects were altered to varying degrees in OSNs of infected patients: As many of these changes involve chromosomes containing a large number of ORs, the altered structure may alter or disrupt key processes dependent on the presence and regulation of key inter-chromosomal contacts.

### Analyzing the relationship between fine structure and regulation of olfactory receptor clusters

Despite limited direct experimental data on OR regulation in human OSNs, functional and structural patterns observed in mice, a well-studied model organism for the olfactory system, prompted us to explore regulation mechanisms in the human samples. Current data on OR regulation in mouse OSNs point to (i) the heterochromatic compaction of OR clusters (26,28), and (ii) activation of selected ORs via looping interactions with enhancers, dubbed the Greek Islands, that are located near corresponding OR clusters (29,31). Understanding how altered chromatin structure affects OR regulation would therefore require us not only to zoom in on a finer genomic scale, but also to consider interactions of ORs with potential enhancers located near OR clusters. We postulated that since many mouse transcription factors have orthologs in humans (39), enhancers in mice may also show some degree of conservation in humans, hence a direct mapping of the Greek Island enhancers found in mice onto the human genome may provide a reasonable set of putative enhancers (PEs) of ORs in human OSNs. We mapped the 35 Greek Islands enhancers from the mouse genome to the human genome, identifying a subset of 15 PEs located within 1 Mb of at least one human OR (mapping results are listed in Supplementary File, Table S1). Noteworthy, PEs considered in this work include: (i) Milos, located <200kbp from the only cluster of Class I ORs located in chromosome 11, (ii) Symi, Lesvos, Skiathos, and H, located on the flanks of two Class II OR clusters in chromosome 14, and (iii) Thira, located within a large Class II OR cluster near the q-arm telomere of chromosome 1. Having obtained the genomic coordinates of ORs and PEs, we then developed an analytical procedure using Hi-C interaction data to identify strongly localized structural units and the interaction energies between them, enabling us to investigate if human ORs are also physically compacted, and if ORs form preferential interactions with the putative enhancers identified above. Figure 3 (see also Supplementary File, Figure S6) shows a schematic diagram illustrating this procedure performed on the genomic region chr11:5.5-6.5Mb: using the Hi-C interaction matrix at 5kb resolution, we began by considering each 5kb bin as an individual structural unit, represented by black circular nodes in the network schematic at the bottom, joined by a curve representing the chromosomal trace. We then iteratively merged pairs of structural units with the highest interaction energy ratios r, defined as the ratio of interaction energy between pairs of distinct units to the average interaction energy within the units. The highest value of *r* in the remaining network, *r_max_* decreases monotonically, and when *r_max_* falls below 1.0 we obtain the set of structural units marked by grey ellipses in the network. A 1D genomic representation of the structural units is shown at the top, where white-colored horizontal bars indicate a structural unit made up of a contiguous series of 5kb bins, while colored bars of the same color indicate non-contiguous segments that join to form a single structural unit at the given level. Decreasing the value of *r_max_* by further merging of units leads to a greater degree of separation between larger structural units (see blue, red, then green ellipses in Figure 3 and Supplementary File, Figure S6, corresponding to the lowest *r_max_* = 0.8,0.6,0.5 (see Materials and Methods), respectively), indicating the presence of a hierarchy in chromatin structure at corresponding scales. In this work, to obtain a sufficient degree of physical separation between structural units we used the cutoff *r_max_* = 0.6. In general, we observed that patient samples had larger interaction patterns within each chromosome, leading to larger structural units compared to control samples (see Supplementary File, Table S2). The structural units obtained for *r_max_* = 0.6 and 0.5 have typical size of (sub)TADs with 100-400 kb sizes, apparently demarcating continuous and discontinuous TADs (37) containing ORs and their regulatory elements. The higher *r_max_* = 0.8. and 1.0 allow to delineate other basic structural units of chromatin, loops (34–36) with characteristic sizes of 30-60kb, forming the (sub)TADs and allowing to explore the structure and interactions in the latter and their alterations between normal and pathological states.

We began by studying the organization of the only cluster of 54 Class I ORs in the region 3.5-6.5Mb of chromosome 11, flanked by a PE Milos and an OSN marker gene CNGA4. Here and below and we perform our analysis on two controls and two patient subjects. Aggregate metrics comparing the sizes and interactions between OR-containing and OR-free units are provided in Table 1. In the interaction heatmaps for each sample (Figure 4), the position and size of structural units are indicated by dark red squares along the diagonal, while strong interactions (with interaction energy ratio *r* > 0.05) between units are marked by colored off-diagonal areas. Adjoined to the interaction heatmaps are two panels that depict the density profile of OR genes (blue plot), while the location of PE Milos and the OSN marker gene CNGA4 are marked by black and red dashed lines, respectively. More structural details are illustrated in the network diagrams (Figure 4), where we represented structural units by circles with radii corresponding to unit sizes: Strongly interacting units are joined by black/grey edges (black edges indicate an interaction energy ratio strong interactions with *r* > 0.5, whereas grey edges indicate interactions with 0.2 < *r* ≤ 0.5, and weaker interactions are omitted for clarity), and the grey curve serves as a chromosomal trace in the 5’-to-3’ direction. The distribution of ORs across structural units is represented by the color saturation of the units, and the location of the PE and the OSN marker gene are represented schematically by the black and red boxes along the chromosomal trace. Beginning with aggregate statistics (Table 1), we observed that in this region, in comparison with control samples, all patient samples formed larger units overall. Also, the average interaction energy ratios *r* between structural units were higher in patient than in controls. The OR-containing units show stronger difference between the patients and controls than OR-free ones (Table 1), pointing to a consistent change in chromatin architecture between controls and patients. At the same time, we observed that given higher or comparable compactness (Table 1), the interactions between OR-containing units were substantially stronger than that of OR-free units in all samples, suggesting that the OR cluster exhibits more compact packing than OR-free regions – consistent with the current view (based on mouse models) of OR compaction and silencing in heterochromatin. Further features of the organization of the OR cluster can be seen in the interaction heatmaps (see Figure 4). In controls there is a clear division of the OR cluster into 3 strongly interacting subsections (approximately 4.2-5.1Mb, 5.1Mb-6.0Mb, and 6.0-6.5Mb) with relatively weak interactions between them. In patients, however, strong long-range interactions within the OR cluster render these subsections barely distinguishable, in agreement with the hypothesis that the OR cluster is more compacted in patients. The network representation of structural units shows further details on the structural environment of key genomic elements: while control samples had the PE Milos located in a small unit free from ORs, in both patient samples we found that the PE Milos formed part of a large structural unit containing multiple ORs (450kb unit with 5 ORs in p102, 300kb unit with 6 ORs in p146). Also, for controls the OSN marker gene CNGA4 (44kb downstream of the nearest OR) resided in OR-free units, but for patients the marker gene was included in the same structural unit as the proximal OR. Taken together with the observation of strengthened OR cluster compaction in patients, and the model of heterochromatic silencing on compacted OR genes in mouse models, the inclusion of the PE Milos and OSN marker CNGA4 in OR-containing structural units may result in the silencing of both genomic elements. Besides the direct impact of CNGA4 downregulation, the inactivation of the PE Milos may also have indirect consequences on OSN function. Given that Class I ORs have been assumed to rely primarily on intra-chromosomal interactions for activation, the silencing of the PE Milos may significantly impair the transcription of Class I ORs if no other enhancers are located within the vicinity - the next closest putative enhancer, P, is located 2.7Mb downstream, more than 600kb from the other end of the OR cluster.

**Figure 4.**
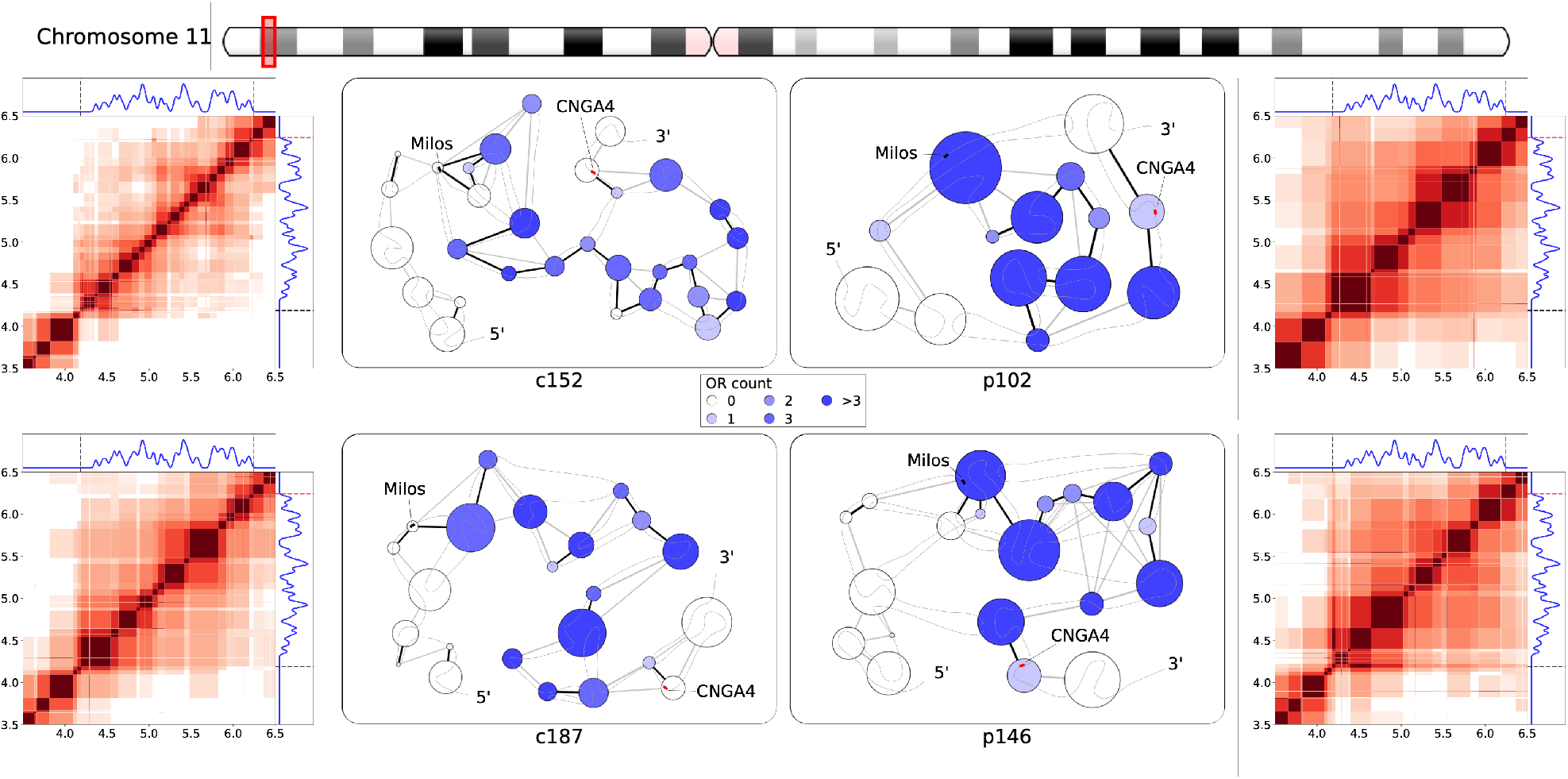
Fine structural organization in the Class I olfactory receptor gene cluster on chromosome 11. Left and right columns. Heatmaps of interactions between structural units in the region 3.5-6.5Mb of chromosome 11. In the heatmaps, dark red squares along the diagonal mark the size and position of continuous structural units, and off-diagonal colored regions indicate strongly-interacting unit pairs (*r* > 0.05). The adjoining panels (above and to the right of the heatmaps) show the 1D distribution of Class I olfactory receptor genes (blue trace), putative enhancers mapped from mouse annotations (black dashed lines), and human mature OSN marker genes (red dashed line). The heatmaps for control samples c152 and c187 are shown on the left column, while patient samples p102 and p146 are shown on the right column. **Center.** Network representations of chromatin structure in the same region 3.5-6.5Mb of chromosome 11, including the PE Milos and the OSN marker CNGA4. Structural units are represented as circles of radii scaling with unit sizes, colored by the number of ORs in the unit. Black/grey edges connecting the units indicate strong interactions, with black edges indicating an interaction energy ratio *r* > 0.5 and grey edges indicate 0.2 < *r* ≤ 0.5. A black trace is overlaid on the network to show genomic positions of the units in the 5’-to-3’ direction on the positive strand. The approximate genomic locations of the PE Milos and the OSN marker CNGA4 are represented by black and red marks respectively.

**Table 1.** Characteristics of structural units identified in neighborhoods of olfactory receptor gene clusters in chromosomes 1, 11, and 14. The average size of structural units and the average interaction energy ratio between structural units in selected regions containing OR clusters, featured in Figures 4–6. The average size and interaction energy ratios for subgroups of OR-containing (ORc) and OR-free (ORf) structural units were also included, along with the ratio of averages for ORc to ORf subgroups.

| Location | OR Class | Sample | Average structural unit size |  |  |  | Average interaction energy ratio |  |  |  |
| --- | --- | --- | --- | --- | --- | --- | --- | --- | --- | --- |
|  |  |  | Overall | ORc | ORf | ORc/ORf | Overall | ORc | ORf | ORc/ORf |
| **Chromosome 11**<br>**3.5-6.5Mb** | I | c152 | 103.7 | 97.9 | 113.6 | 0.86 | 0.115 | 0.164 | 0.108 | 1.52 |
|  |  | c187 | 144.8 | 147.1 | 141.1 | 1.04 | 0.125 | 0.197 | 0.150 | 1.31 |
|  |  | p102 | 235.4 | 206.4 | 341.7 | 0.60 | 0.197 | 0.254 | 0.178 | 1.43 |
|  |  | p146 | 165.0 | 170.4 | 156.9 | 1.09 | 0.153 | 0.248 | 0.170 | 1.46 |
| **Chromosome 1**<br>**246-249Mb** | II | c152 | 101.0 | 80.3 | 135.0 | 0.59 | 0.093 | 0.150 | 0.138 | 1.09 |
|  |  | c187 | 94.7 | 82.6 | 107.5 | 0.77 | 0.107 | 0.168 | 0.164 | 1.02 |
|  |  | p102 | 163.9 | 119.6 | 212.3 | 0.56 | 0.131 | 0.207 | 0.184 | 1.13 |
|  |  | p146 | 141.7 | 98.9 | 191.7 | 0.52 | 0.121 | 0.193 | 0.119 | 1.62 |
| **Chromosome 14**<br>**19.5-22Mb** | II | c152 | 95.4 | 72.1 | 112.8 | 0.64 | 0.091 | 0.133 | 0.107 | 1.24 |
|  |  | c187 | 103.5 | 86.4 | 118.1 | 0.73 | 0.100 | 0.118 | 0.093 | 1.27 |
|  |  | p102 | 134.2 | 131.1 | 137.0 | 0.96 | 0.127 | 0.175 | 0.101 | 1.73 |
|  |  | p146 | 121.1 | 146.8 | 95.5 | 1.54 | 0.098 | 0.158 | 0.090 | 1.76 |

Turning to the Class II ORs scattered across the human genome, we observed that these OR clusters show rather consistent trends in structural organization, which is exemplified here with the large cluster of 43 Class II ORs 246.0-248.9Mb in chromosome 1. Table 1 indicates that OR-containing structural units are smaller in size (0.52-0.77) than OR-free units in this region. Interactions in OR-containing units are stronger than that in OR-free units, and patient samples show a greater enrichment of interactions in general. Figure 5 shows the interaction heatmaps and network representations of structure, in the distribution of ORs denoted by the green density plot and color saturation in structural units and a single PE Thira identified in this region. Comparing the interaction heatmaps for controls and patients, we observed firstly that the region upstream of the OR cluster (approximately 246.0-247.5Mb) showed little interaction with the OR cluster in controls, but in patients this region tends to interact more strongly with the whole OR cluster. Also, the telomeric region (248.8-248.9Mb) interacts with both ends of the OR cluster more strongly in patients (with *r* ≈ 0.10) than in controls (*r* < 0.05). The network representations illustrates that the final ~300kb of the OR cluster (248.4-248.7Mb) forms a set of structural units comprising interspersed genomic segments, indicating a high degree of complexity in chromatin organization in that region. In the region 247.5-248.4Mb, control samples yield a series of weakly-interacting, small structural units, whereas in patients a series of small but strongly interacting units this region (observed in p102) or large units with weak long-range interactions between them (observed in p146). In comparison, control samples adopt a relatively sparse conformation in this section, forming a series of small units interact strongly only with genomically adjacent units. This picture indicates a greater degree of physical compaction of the OR cluster in patients compared to controls, in agreement with the observations made about the Class I OR cluster. Together with the stronger interactions between the OR cluster and the telomeric region, above observations may suggest that the increased physical compaction of this region in patients may be linked to the “spread” of telomeric heterochromatin. Surprisingly, the only regulatory element located in this region, the PE Thira, was located within OR-containing units in both the controls and patients. While physical compaction of the OR cluster may lead to inactivation of this PE, work on mouse models indicate that inter-chromosomal enhancer-promoter interactions play a key role in regulating Class II ORs (31), potentially availing other PEs for OR regulation in this cluster. As the available Hi-C dataset is insufficient for studying inter-chromosomal interactions robustly at this resolution, further work is required to investigate how changes in inter-chromosomal interactions may affect regulation of Class II ORs. Further assays would also be required to identify regulatory elements human ORs to elucidate epigenetic mechanisms of Class II ORs in normal and infected human OSNs.

**Figure 5.**
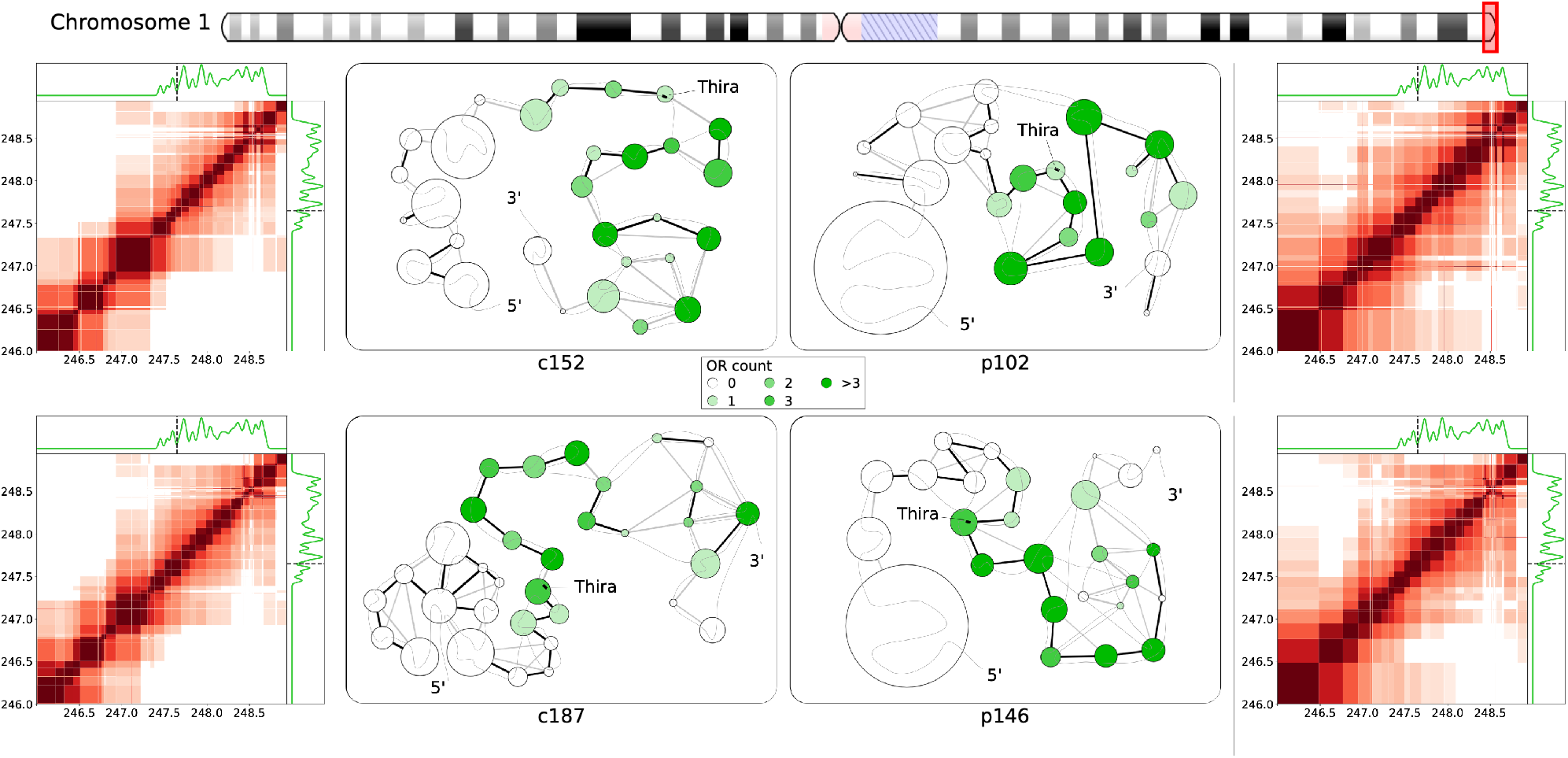
Fine structural organization in a Class II olfactory receptor gene cluster near the chromosome 1 q-ter telomere. Notations and marks are as in Figure 4. Heatmaps of interactions and network representation are given for structural units in the 246.00-248.94Mb region of chromosome 1. Structural units are indicated by dark red squares along the diagonal, and off-diagonal colored regions show interactions with energy ratios above a threshold, *r* > 0.05. The distribution of Class II OR genes is indicated by the adjoining 1D plots by the green traces, and the location of the PE Thira is marked by the black dashed line. Network representation of structural units in the OR cluster, with strong interactions indicated by black/grey edges, and the approximate location of the PE marked by a black mark.

A further example of Class II OR clusters can be found in the pericentromeric region of chromosome 14, around the coordinates 19.5-22.0Mb, with 4 PEs mapped into the region. This region comprises two separate clusters, with a larger one (19.8-20.3Mb) containing 17 ORs, flanked by the PEs Symi and Lesvos, and a smaller one (21.5-21.7Mb) containing 3 ORs, flanked by PEs H and Skiathos. Table 1 shows that patients reveal larger structural units than controls as well as stronger interactions in them, which is chiefly result of stronger interactions in OR-containing units. Figure 6 provides further details, with the interaction heatmap zoomed out to include the pericentromeric region from 18.0Mb. The green frame marks out the region 19.5-22.0Mb that we focus on in the network representation, where we included a red node to represent the large, multi-segment unit associated with centromeric heterochromatin. The interaction heatmaps for controls show a separation of the upstream cluster from the downstream OR-free region, while in patients we observed stronger interactions between the two. The OR cluster also tends to form sparser interactions in controls than in patients, concurring with our general observations in Table 1. The network representation shows that the upstream OR cluster in controls formed only a series of small units in a loosely packed conformation, yielding strong interactions only with genomically adjacent units (despite small segments grouped with centromeric heterochromatin in c187) similar to structural units and interactions between them in controls of the Class II OR cluster in Figure 5. In patient samples the upstream cluster formed a set of highly interlinked units with centromeric heterochromatin, while downstream region reveals formation of larger structural units, much like that previously observed in chr1:248.4-248.7Mb. Shifting our focus to PEs, while Symi and Lesvos were consistently located in OR-containing units for all samples, there is a movement of H and Skiathos between OR-containing/free units with no consistent difference observed between patients and controls. Notably, all PEs are observed in larger structural units in patients compared to controls.

**Figure 6.**
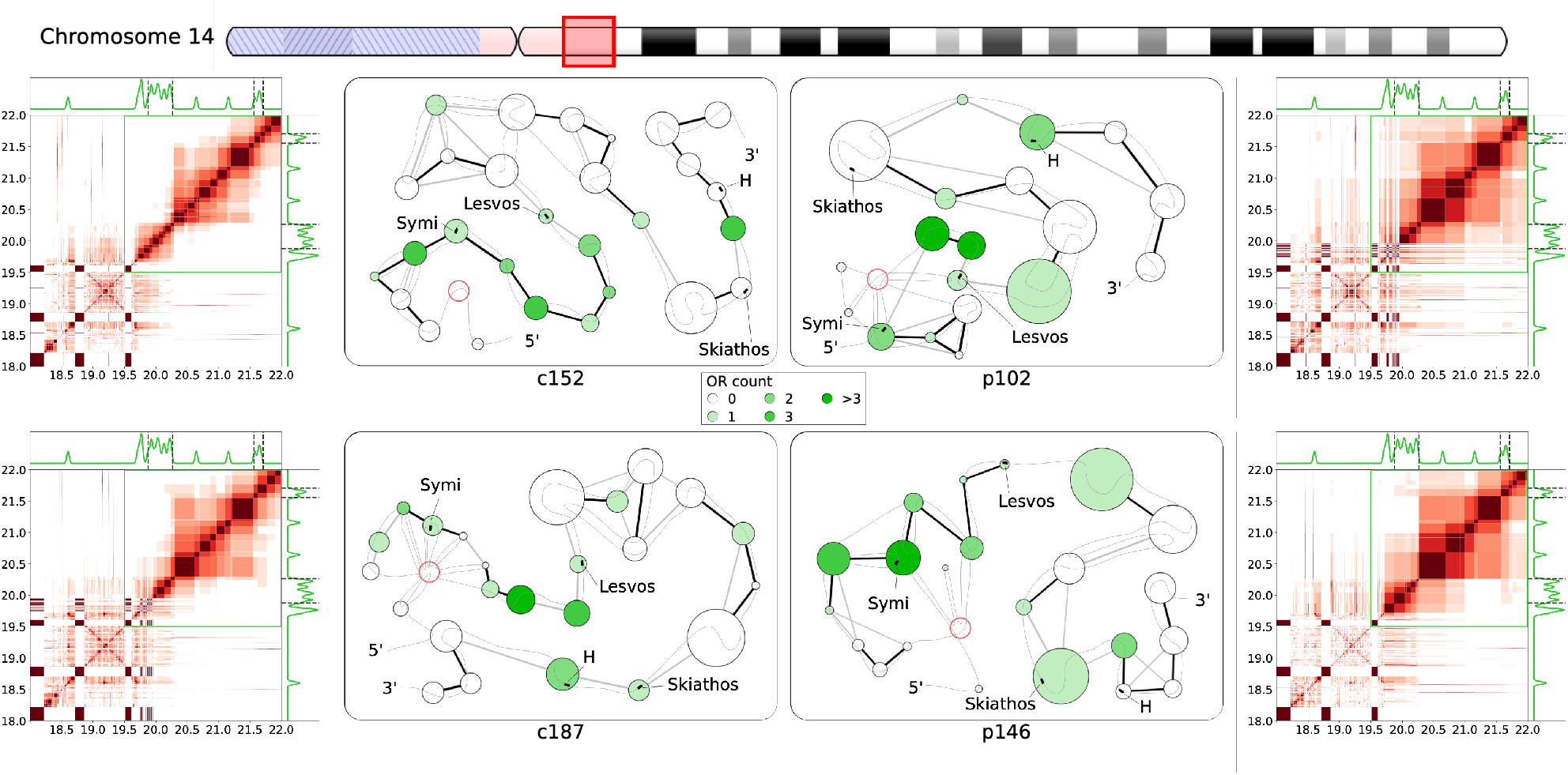
Fine structural organization in two Class II olfactory receptor gene clusters near the chromosome 14 centromere. Notations and marks are as in Figure 4. Heatmaps of interactions and network representation are given for structural units in the region 18.0-22.0Mb of chromosome 14. Two OR clusters with nearby PEs are present in this region: the cluster at 19.8-20.3Mb contain PEs Symi and Lesvos, while the smaller cluster 21.5-21.7Mb is flanked by PEs Skiathos and H. In the network representation the large structural unit associated with pericentromeric heterochromatin is composed of multiple non-contiguous segments, and is represented here by the single node with a red border.

Summarizing the analysis of the internal organization of OR gene clusters by identifying fine structural units using high-resolution Hi-C data and by quantifying the interaction energy between these units, we obtained the following general observations: In most of the cases the average size of OR-containing structural units is smaller than that of OR-free ones. The OR-containing units also form consistently stronger mutual interactions than OR-free counterparts in the same regions, with a stronger bias seen for patients in Class II OR clusters (Table 1). Patient samples generally form larger structural units and stronger interactions. The interactions in OR-containing units are always stronger than in OR-free ones in both controls and patients with more pronounced effect in the latter. Network representations of structural units in OR clusters illustrate less compact, more linear organization of OR-containing units in controls, while patients either formed larger units overall, or formed a series of small units with complex interlinking or strong interactions between them (Figures 4–6). These observations suggest stronger physical compaction and potential involvement in silencing interactions of OR clusters in pathology, which can disrupt OR expression leading to anosmia in COVID-19 patients. The increased structural units and their compaction can be a general mechanism of OR silencing, and the grouping of the upstream PE Milos and the downstream OSN marker gene CNGA4 in the OR-containing units in patients in the Class I OR cluster (Figure 4) may work as a complementary mechanism by which olfaction may be impaired in COVID-19 infected patients.

## Discussion

While anosmia has been a useful diagnostic indicator for early-stage COVID-19 infection (1–4), mechanisms behind its rapid onset and recovery has been a subject of debate. As human olfactory sensory neurons (OSNs) do not express the primary receptors that enable direct infection by the SARS-CoV-2 virus, it has been proposed that inflammatory response and damage to support cells in the olfactory epithelium (OE) may result in olfactory impairment (8–11). Furthermore, as most patients recover their sense of smell within a short time frame of 1-2 weeks (12–15) it is unlikely that these changes have led to massive cell death among OSN populations, thus we set out to understand how COVID-19 infection in the OE may affect olfactory signal transduction in OSNs non-cell autonomously. Given that olfactory receptors (ORs) are vital in initiating olfactory signal transduction in OSNs, a recent study linking inflammatory cytokine signaling to decreased OR expression (22), in particular, upregulation of certain interferon regulated genes leading to chromatin reorganization as a result of the immune response, prompted us to investigate how OR regulation is determined and modulated by the chromatin organization. Considering earlier observed specifics of OR regulation in the mice chromatin structures (27, 28, 31), we hypothesized here that the disruption of the OR function can be caused by chromatin reorganization taking place upon SARS-CoV-2 infection. Specifically, we used Hi-C data on human OSNs to study if features identified in mouse models, such as the interchromosomal aggregation of OR genes (27, 38) and looping interactions between OR clusters and nearby enhancers (29), have identifiable parallels in humans, and if these features are disrupted upon COVID-19 infection in the OE.

To that end, we first reconstructed whole-genome chromatin ensembles using a stochastic embedding procedure we have previously developed (32). While the relative positioning of chromosomal territories (CTs) in SARS-CoV-2-infected patients and control subjects were somewhat similar (see Figure 1), we observed that chromosomes containing the most OR genes tend to intermingle strongly, consistent with the interchromosomal OR aggregation observed in mice (28). This intermingling was significantly weakened in infected patients: notably, chromosomes 1 and 9 showed significant detachment from all other chromosomes in patient samples. While having chromosomal intermingling cut off may hinder the formation of interchromosomal enhancer hubs (30, 31), understanding how structural features of chromatin relate to OR regulation required us to probe finer levels of detail, looking at chromatin architecture at the level of OR gene clusters and putative enhancers (PE) mapped from mouse data. To this end, we developed an analytical method to identify local structural units down to an average size of (sub)TADs, 100-200kb and further down to forming TADs chromatin loops with characteristic sizes of 30-60kb in order to explore structural units and interactions between them relevant to OR regulation. We observed that OR-containing structural units had consistently stronger mutual interactions than OR-free units in all samples — indicative of the heterochromatic physical compaction of OR genes observed experimentally (27, 28) — though this effect was more pronounced for patients in Class II OR clusters (Table 1). The stronger interactions observed between OR-containing units leading to formation of larger structural units are more pronounced in patients (Figures 4–6), pointing to a potential disruption of chromatin organization within OR clusters for COVID-19 patients. Further, we observed that key genomic elements near the Class I OR cluster (Figure 4), including the only PE (Milos) in that region and a marker gene critical for OSN function (CNGA4), were grouped into OR-containing structural units in patients: the heterochromatic compaction of ORs may lead to these genomic elements being inactivated in patients, potentially also leading to olfactory dysregulation. The structural context of ORs and their regulatory elements in Class II OR cluster will have to be further studies upon availability of corresponding data. Overall, despite many open questions about functional significance of structural details in OR clusters due to insufficient data, our analysis here indicates significant alterations on both inter- and intrachromosomal levels in the chromatin of the COVID-19 patients. Specifically, the structural environment of OR-rich chromosomes investigated at the level of chromosomal territories and their intermingling reveals an alternative conformation providing interactions between OR-containing chromosomes associated with SARS-CoV-2 infection. The local intrachromosomal structural hierarchy in locations of ORs genes, their enhancers and other epigenetic factors yields formation of (sub)TADs from chromosomal loops and alteration of their structure in pathological samples that may contribute to the onset of anosmia in COVID-19 patients. Given the importance of understanding molecular mechanisms of SARS-CoV-2 infection for future development of sensitive diagnostics and drugs for emerging new variants of concern (VOCs), the genomic and epigenomic studies should be further extended and developed into more precise approaches with predictive and design capabilities (40). These tasks will require a number of improvements in resolution of experimental techniques and additional data on epigenetic factors and their mapping on genomic sequences: (i) it should include further development of the Hi-C analysis and relevant techniques (41–43) for mapping epigenetic signals, their merging in combined pipelines (44,45), and consideration of chromatin-lamina interactions in the reconstruction of the 3D chromatin structure, to name a few; (ii) currently missing information on relevant regulatory elements and epigenetic signals (46,47) should be also obtained and mapped on genomic sequences for obtaining a complete picture of the genome expression and regulation in norm and pathology.

## Supporting information

MS_SUbmission_BA

