## Supplementary material for "Disrupted chromatin architecture in olfactory sensory neurons: A missing link from COVID-19 infection to anosmia": MS_SUbmission_BA

| Enhancer name | Mouse genome coordinates (mm9) | | | Human genome coordinates (hg38) | | | Near human OR gene? |
|---|---|---|---|---|---|---|---|
| | Chr | Start | End | Chr | Start | End | |
| **EVIA** | 2 | 111566700 | 111567700 | 1 | 421646 | 422646 | Y |
| **THIRA** | 7 | 93443100 | 93444300 | 1 | 247647764 | 247648964 | Y |
| **GAVDOS** | 16 | 58956100 | 58957500 | 3 | 98333106 | 98334506 | Y |
| **LIPSI** | 2 | 36982900 | 36984500 | 9 | 122693320 | 122694920 | Y |
| **KALYMNOS** | 6 | 116563100 | 116564900 | 10 | 45049357 | 45051157 | Y |
| **MILOS** | 7 | 109661022 | 109662409 | 11 | 4186792 | 4188179 | Y |
| **P** | 7 | 114190700 | 114192100 | 11 | 6823977 | 6825377 | Y |
| **IKARIA** | 7 | 106936000 | 106937199 | 11 | 75035882 | 75037081 | Y |
| **SYMI** | 11 | 50812200 | 50813599 | 14 | 19872923 | 19874322 | Y |
| **LESVOS** | 14 | 51377700 | 51379500 | 14 | 20261516 | 20263316 | Y |
| **SKIATHOS** | 14 | 52957700 | 52959300 | 14 | 21556796 | 21558396 | Y |
| **H** | 14 | 53166900 | 53168700 | 14 | 21703296 | 21705096 | Y |
| **CRETE** | 11 | 73848400 | 73850199 | 17 | 3317487 | 3319286 | Y |
| **SIFNOS** | 11 | 87749900 | 87751500 | 17 | 58133191 | 58134791 | Y |
| **SIKINOS** | 9 | 18525100 | 18526900 | 19 | 9054150 | 9055950 | Y |
| **NIMOS** | 4 | 118314100 | 118315900 | 1 | 43129059 | 43130859 | N |
| **KOS** | 3 | 106678500 | 106680100 | 1 | 110797728 | 110799328 | N |
| **FOLEGANDROS** | 3 | 97294700 | 97296500 | 1 | 147359887 | 147361687 | N |
| **PONTIKOS** | 3 | 97141900 | 97143500 | 1 | 147494604 | 147496204 | N |
| **SKORPIOS** | 3 | 97113500 | 97114900 | 1 | 147527207 | 147528607 | N |
| **OTHONI** | 3 | 97111500 | 97113500 | 1 | 147528471 | 147530471 | N |
| **CORFU** | 19 | 14321300 | 14322900 | 9 | 79969332 | 79970932 | N |
| **ALONISOS** | 19 | 14314300 | 14315900 | 9 | 79976205 | 79977805 | N |
| **AKTHRA** | 19 | 14300700 | 14301900 | 9 | 79983966 | 79985166 | N |
| **TINOS** | 19 | 14170300 | 14171700 | 9 | 80089752 | 80091152 | N |
| **RHODES** | 1 | 94441500 | 94443500 | | | | |
| **HYDRA** | 2 | 112049700 | 112051300 | | | | |
| **NISYROS** | 3 | 106683500 | 106684900 | | | | |
| **SFAKTIRIA** | 6 | 42818900 | 42820700 | | | | |
| **KEFALLONIA** | 7 | 6497300 | 6499100 | | | | |
| **IOS** | 7 | 115939700 | 115941900 | | | | |
| **FOURNI** | 7 | 147372100 | 147374100 | | | | |
| **LEFKADA** | 11 | 52014700 | 52015900 | | | | |
| **KARPATHOS** | 14 | 50681500 | 50682900 | | | | |
| **LEMNOS** | 15 | 98340700 | 98342300 | | | | |

**Supplementary File, Table S1. Mapping of Greek Islands enhancers from the mouse genome to the human genome.** List of Greek Islands enhancers identified on the mm9 mouse reference genome, with genomic coordinates listed. Enhancers that have corresponding regions in the human reference genome hg38 have genomic coordinates mapped using the UCSC LiftOver tool. A mapped enhancer located near (within 1Mb from) at least one OR gene is considered as a putative enhancer in this work.

| Chromosome | Sample | Energy ratio cutoff | | | |
|---|---|---|---|---|---|
| | | **1.0** | **0.8** | **0.6** | **0.5** |
| **1** | c152 | 32.5 | 48.5 | 108.8 | 218.7 |
| | c187 | 30.4 | 46.1 | 109.5 | 246.8 |
| | p102 | 31.7 | 53.3 | 184.8 | 370.4 |
| | p146 | 28.4 | 44.0 | 129.2 | 292.1 |
| **11** | c152 | 32.9 | 49.4 | 111.7 | 220.8 |
| | c187 | 30.8 | 46.3 | 115.9 | 254.5 |
| | p102 | 32.5 | 56.4 | 199.4 | 446.1 |
| | p146 | 28.4 | 44.2 | 138.0 | 353.7 |
| **14** | c152 | 32.6 | 48.6 | 104.1 | 204.7 |
| | c187 | 30.9 | 46.0 | 108.7 | 246.8 |
| | p102 | 32.2 | 55.3 | 181.4 | 407.1 |
| | p146 | 28.3 | 44.8 | 127.7 | 318.3 |

**Supplementary File, Table S2. Average structural unit sizes at different levels of hierarchy.** For each sample, we identified structural units at the whole-chromosome level with interaction energy ratios down to various cutoff values, $r_{max} = 1.0, 0.8, 0.6, 0.5$ (see also Materials and Methods for further explanations). The corresponding average size of structural units for each case is listed.

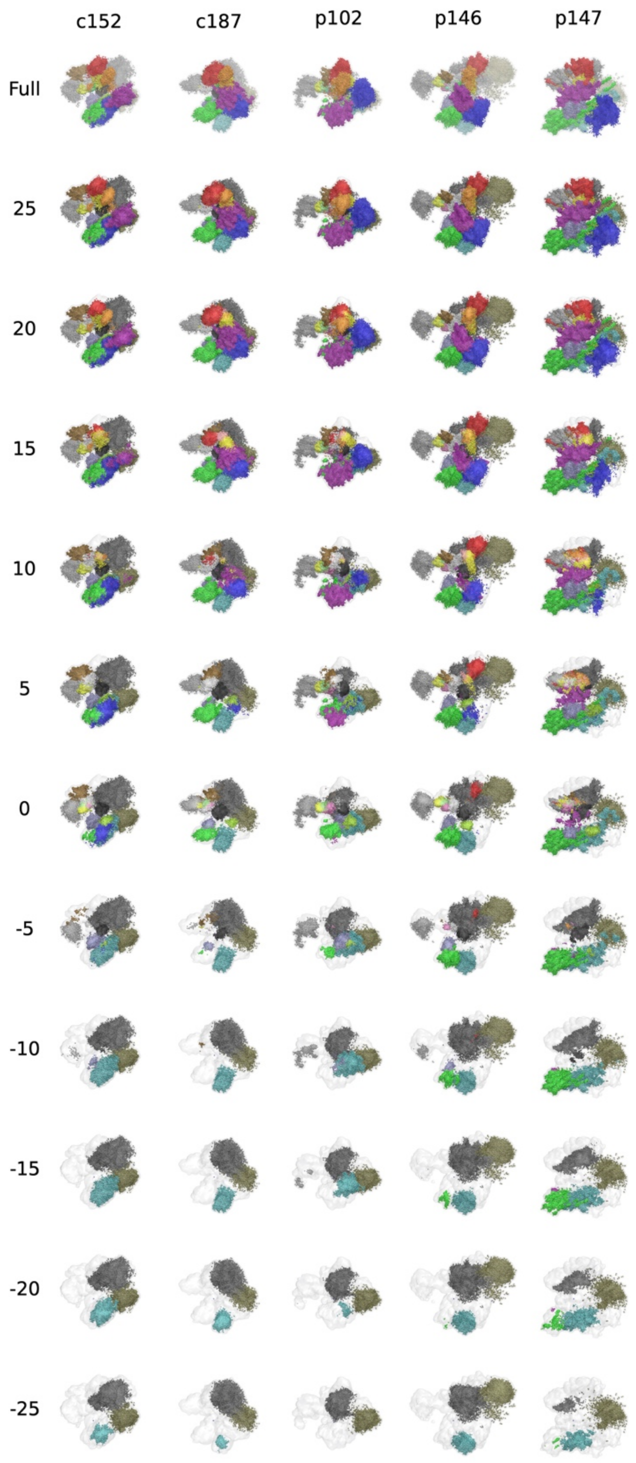


**Supplementary File, Figure S1. Cross-sectional views of OSN chromatin ensemble reconstructions**. Cross-sectional views of chromatin ensemble reconstructions, from $z = +25$ a.u. to $z = -25$ a.u.: corresponding values of $z$ are indicated on the left of each row.

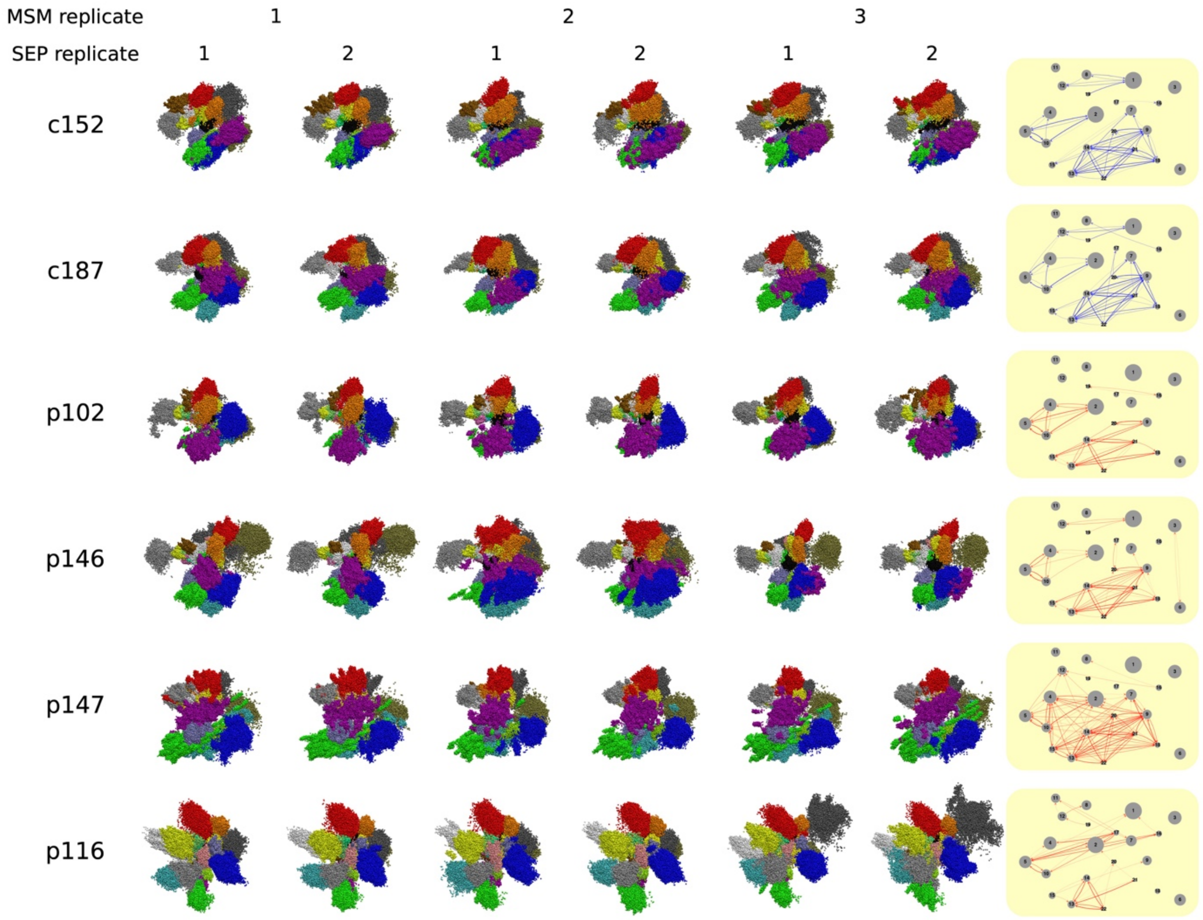


**Supplementary File, Figure S1. Comparing MSM- and SEP-replicates of whole-genome reconstruction on patient and control OSN Hi-C data. Left.** For each Hi-C sample, we performed 3 replicates of MSM-based partitioning to identify structural units, and in each case performed 2 replicates of SEP ensemble reconstructions. The ensemble structures are shown, colored by chromosomes. **Right.** Network representation of chromosomal intermingling in each Hi-C sample. Chromosome pairs with strong intermingling on average (across 6 reconstruction replicates) are joined by an edge, colored blue/red in control/patient samples.

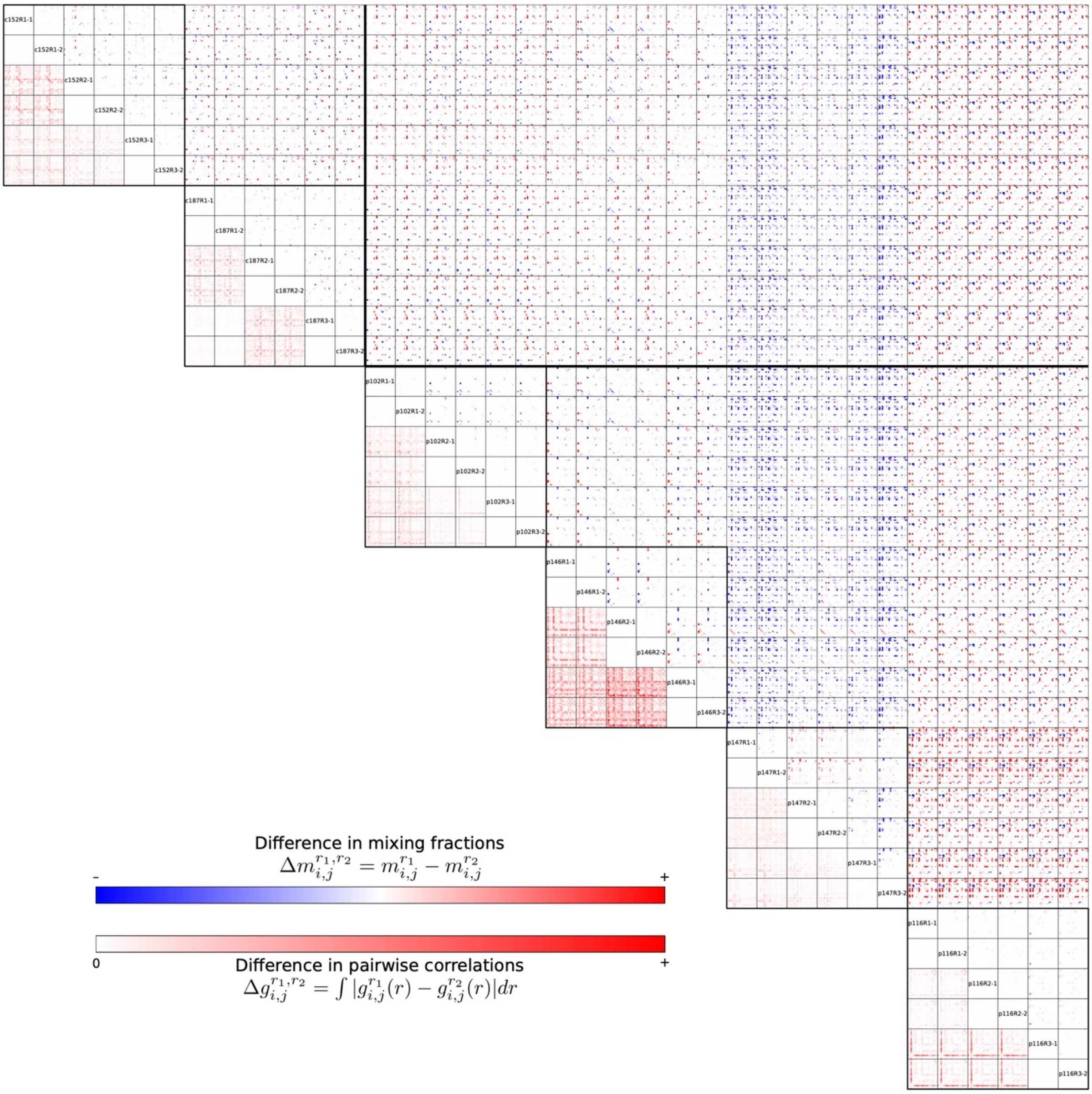


**Supplementary File, Figure S3. Comparing structural metrics shows robustness of reconstructions.** The metrics of chromatin structure between MSM and SEP replicates of the same Hi-C sample, and between Hi-C samples were compared. On the lower left side of the diagonal, comparison of reconstruction replicates on the same Hi-C sample by plotting the heatmap of histogram differences in pairwise correlation functions $\Delta g_{ij}^{r_1,r_2}$ is presented. On the upper right side of the diagonal, the differences in mixing fractions $\Delta m_{ij}^{r_1,r_2}$ are plotted.

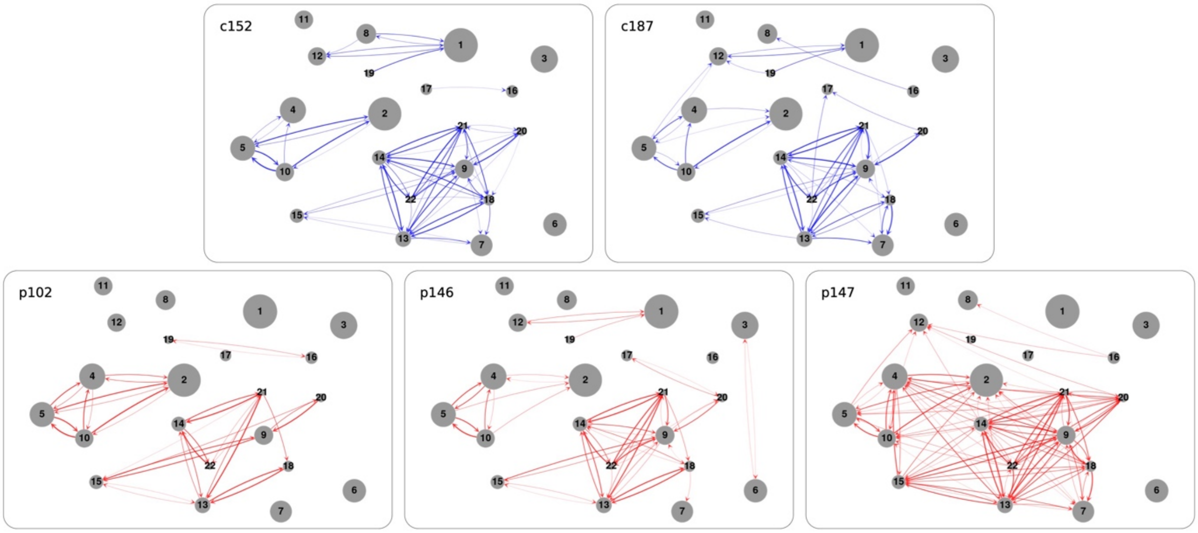


**Supplementary File, Figure S4. Chromosomal intermingling network for all Hi-C samples.** Network of strongly intermingling chromosomal territories in each sample. For each sample, edges connecting CTs $i \rightarrow j$ indicate that more than 10% of chromosome $i$ is located within 1 a.u. from chromosome $j$.

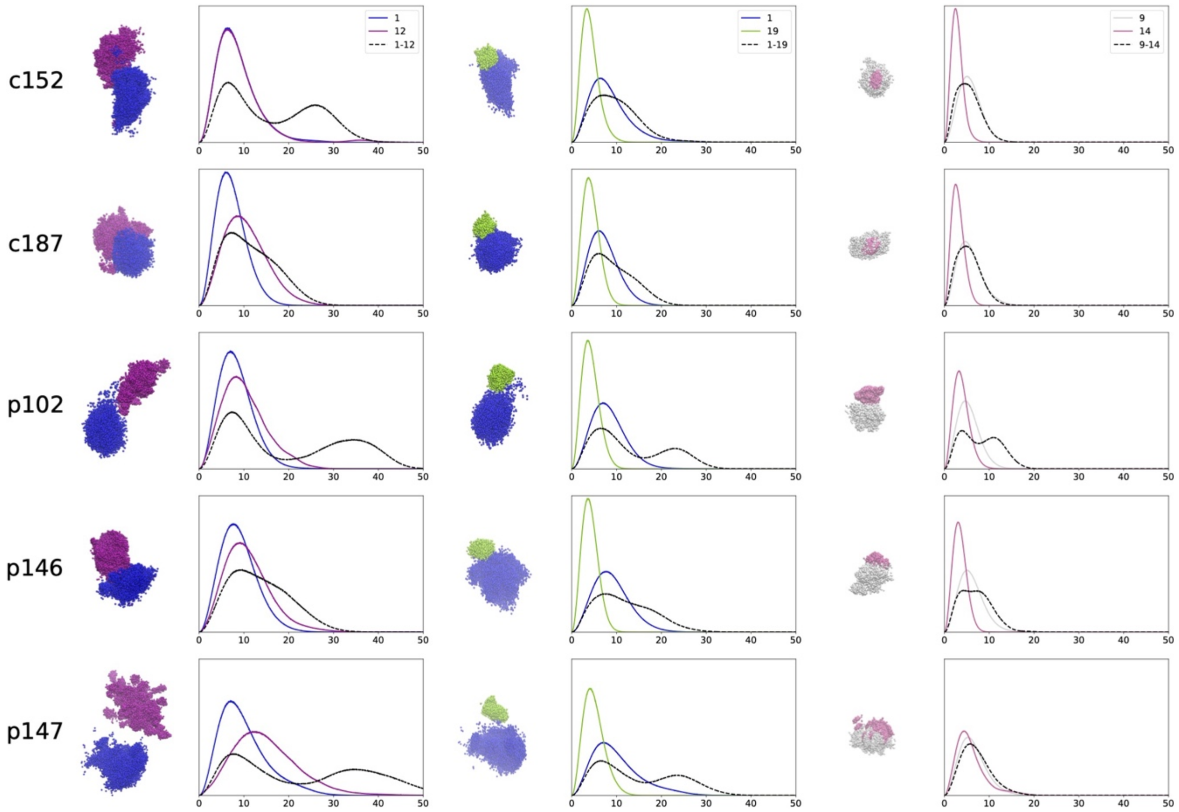


**Supplementary File, Figure S5. Chromosomal territory morphology and distance correlation plots for sample CT pairs for all Hi-C samples).** Structures and $g(r)$ plots are shown here for all samples.

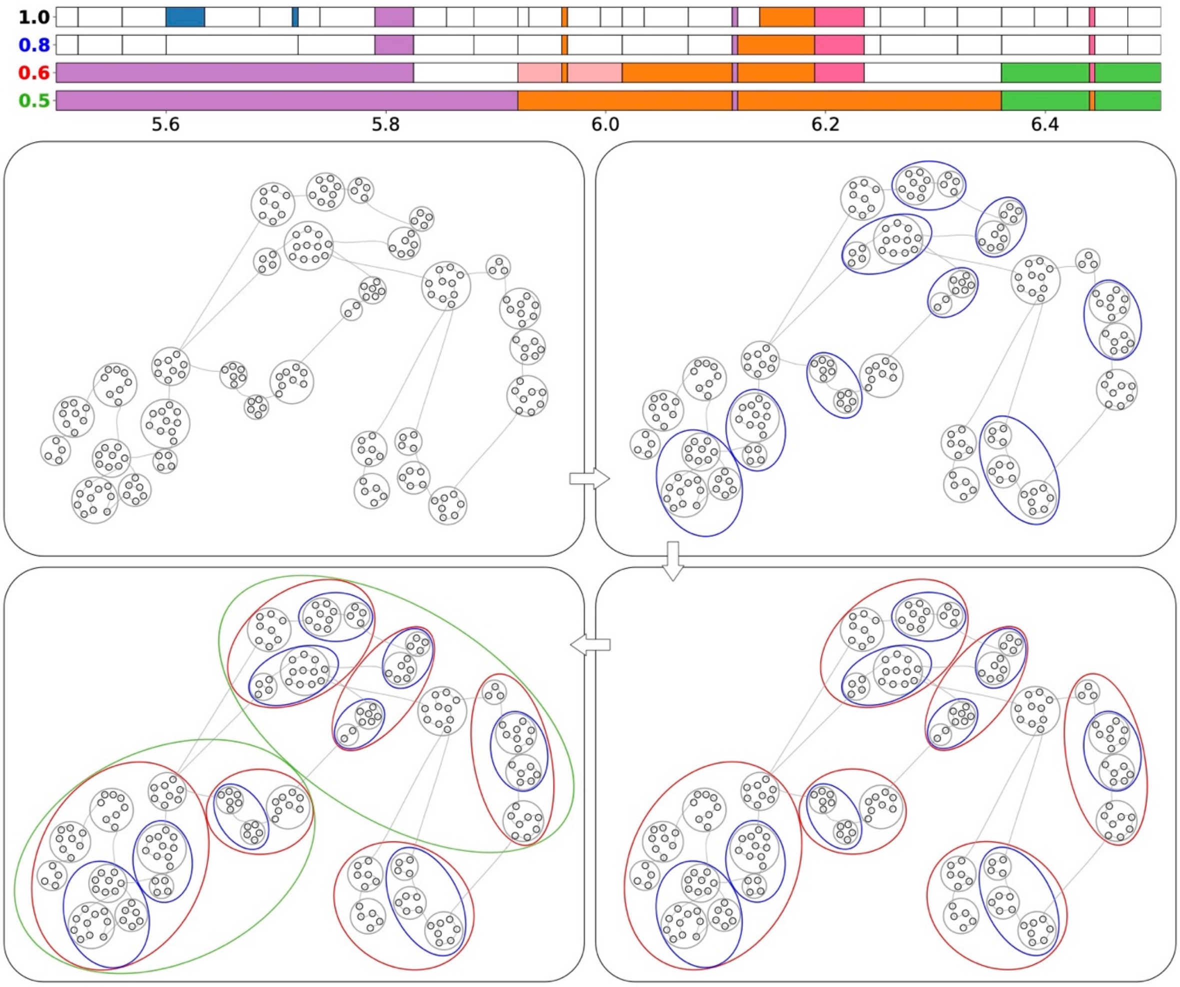


**Supplementary File, Figure S6. Iterative procedure for identifying local hierarchy of the fine structural units.** This Supplementary File, Figure Shows Structural units identified at various levels of hierarchy in a 1Mb region of chromosome 11 (5.5-6.5Mb) are shown here. Individual 5kb bins/loci in this region are represented by black circular nodes, with a chromosomal trace overlaid to show the relative genomic locations of each locus. Beginning by considering each locus as an individual structural unit, we iteratively join pairs of structural units with the highest interacting energy ratios $r$, to a point where the largest energy ratio $r_{max}$ falls below 1.0, yielding structural units denoted by the grey ellipses, in the first panel on the top left. Continuing with this procedure, we obtain a deeper level of hierarchy when $r_{max}$ falls below 0.8, where some of the grey units are merged into larger units marked by the blue ellipses in the second panel to the right. This iterative procedure can be applied further to obtain better-separated structural units with lower values of $r_{max} = 0.6, 0.5$, merging to form larger units marked by red and green ellipses respectively. The 1D representations of the structural units (shown in the top) obtained at each level of hierarchy are denoted by the value of $r_{max}$ on the left. Units composed of a single contiguous genomic segment are shown as white horizontal bars in the plot, while units composed of non-contiguous segments are color-coded accordingly: *e.g.,* the two blue segments and the two orange segments at $r_{max} = 1.0$ form two corresponding structural units.
